# The creation of ghost forests is driven by physical, ecological, and disturbance factors but remains a rare phenomenon

**DOI:** 10.1101/2021.09.14.460301

**Authors:** Christopher R. Field

## Abstract

Marine transgression and the landward migration of coastlines in response to sea-level rise will determine the future of coastal ecosystems and species worldwide, yet no complete model of these processes exist. Ghost forests, areas where coastal forest has been killed by saltwater inundation, are an attention-grabbing early indicator of marine transgression. Research on marine transgression to date has largely been limited to place-based studies that, due to logistical constraints and foci of expertise, investigate subsets of the potential drivers of marine transgression. Here I take advantage of new open source datasets to provide the first systematic analysis of marine transgression across large scales, focusing on the creation of ghost forests across a region of ecological and scientific importance, the northeast U.S. My analysis provides the first synthesis of the physical, ecological, and disturbance factors that influence marine transgression. It also provides crucial regional context for more intensive studies that have a limited geographic scope. I found that ghost forests are a rare phenomenon in the landscape context: 95% of recent forest loss is concentrated in less than 3.86% of marshes, and between 2000 and 2018, only 0.88% of the entire forested area of the transgression zone experienced loss. As a result of this rarity, regional variation in marine transgression is largely driven by opportunities for rare events, which are more numerous when suitable conditions, such as shallow slopes, cover extensive areas. Quantifying recent trends in the rate of forest loss found little evidence of acceleration, with evidence instead suggesting that fewer ghost forests are being created. I also found that physical, ecological, and disturbance factors, including hurricane impacts, were all important for understanding recent trends in forest loss, suggesting that an interdisciplinary approach is warranted for future analyses and modeling of marine transgression. Such interdisciplinary research is urgently needed, as the current rate of marine transgression points to the likelihood of near-term losses of coastal wetlands, with dire implications for the species that depend on them.

## Introduction

Marine transgression, the process by which shorelines move to higher ground in response to sea-level rise, is causing the conversion of upland to wetland ecosystems worldwide (Kirwan & Gedan 2019). This process is likely necessary to prevent losses of coastal wetlands (Kirwan *et al.* 2016), and will have significant economic consequences for coastal cities and communities (Bin & Polasky 2005; Haer *et al.* 2013), which often act to prevent further transgression (Field *et al.* 2017a). Given the potentially steep trade-offs associated with ecosystem transition, there is widespread interest among both researchers and practitioners in understanding the physical and ecological factors that drive, and could potentially be used to predict, the rate and extent of marine transgression and shoreline conversion (Kirwan & Gedan 2019).

Individual studies have investigated subsets of these physical and ecological factors, but they often have limited spatial representation due to the constraints of field studies, which can lead to potentially contradictory results that obscure a general understanding of marine transgression. For example, previous studies suggest substantial variation in the rate of marine transgression, raising questions about which rates are representative of the larger regional trend (Wasson *et al.* 2013; Field *et al.* 2016; Kirwan & Gedan 2019). Despite recent progress in understanding marine transgression (Kirwan & Gedan 2019), key knowledge gaps remain including basic information such as the rate of transgression, spatial and temporal trends, especially the potential for acceleration, and whether current rates will offset losses of coastal wetlands. This information is foundational for coastal management and planning (ACJV 2020) as well as research efforts to quantify the future of tidal marsh ecosystems, including the species that depend on them (Hodgman *et al.* 2015).

Another key gap in our understanding of marine transgression is which physical and ecological factors influence the process, including the relative strength of the most important factors. This information is needed to inform efforts to predict where and when marine transgression is likely to happen, which is critical for developing realistic projection models of tidal marsh extent. These models, often referred to as sea-level rise impact models, have received widespread attention (e.g. the Sea Levels Affecting Marshes Model [SLAMM]; [Wu *et al.* 2015]), but still lack key components that would make them suitable for informing conservation and policy (Kirwan & Guntenspergen 2009).

Here I present the first comprehensive analysis of marine transgression over large scales in which I quantify 1) temporal and spatial trends in rates of transgression, including any evidence for acceleration that might signal the potential for tipping points, and 2) the factors that influence marine transgression, including physical, ecological, landscape, and disturbance-related factors, which have not yet been investigated in the same analytical framework. I used as an indicator of marine transgression the creation of ghost forests, areas where coastal trees have died because of saltwater inundation. Ghost forests, in addition to being the most striking indicator of marine transgression (Kirwan & Gedan 2019), can be detected by remote sensing, making it possible to quantify the factors that influence the creation of ghost forests across areas large enough to capture regional differences that might be limiting inference from smaller studies.

I identified the potential predictors of coastal forest loss based on previous research on the drivers of marine transgression and the availability of large-scale datasets (see **Methods**): the slope of the upland, rate of sea-level rise, proportion of upland that is covered by conifers, height of the peak storm surge during 2012’s Hurricane Sandy, geomorphic setting, and size of the forest patch. I investigated the relative importance of these factors for explaining the rate of coastal forest loss using a Bayesian hierarchical approach that quantified the relative influence of these factors on both the presence and extent of forest loss. I used two approaches to increase the confidence in the results from these correlational analyses: First, for each potential predictor, I made an *a priori* prediction for the direction of the effect based on previous research (cf. O’Connor *et al.* 2015). Second, I developed a simulation to test the model structure and ensure that the potential influence of the independent variables could be distinguished from each other and other potential confounding variables. I applied these methods to forest loss data from the entire marine transgression zone of the northeastern U.S. between 2000-2018 (Figures 1, 2), as this study area is the focus of ongoing research and conservation focused on the responses of coastal ecosystems to sea-level rise (e.g. Lentz *et al.* 2016). Together, this new knowledge on the rates of recent marine transgression over larges scales – coupled with the first estimates of the relative importance of physical, ecological, geomorphic, and disturbance factors – advances the current knowledge of processes underlying marine transgression. This new knowledge establishes the mechanisms and patterns that should form the basis of future projection modeling and highlights the potentially dire challenges facing coastal ecosystems, especially tidal marshes, in the face of sea-level rise.

**Figure 1.**
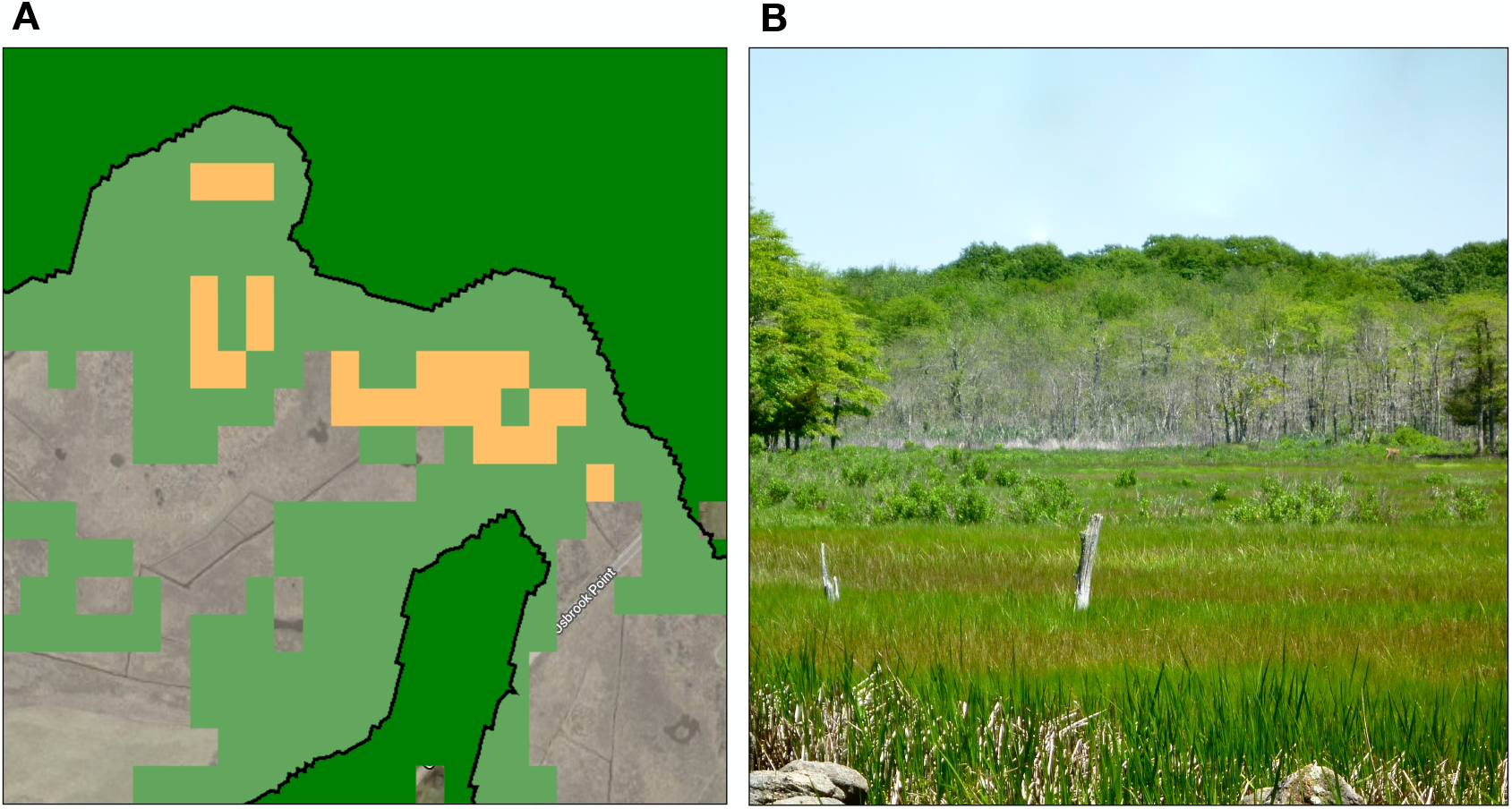
Example of a ghost forest from Stonington, Connecticut, U.S. **A** shows detection of the forest loss, shown in the picture in **B**, using remotely sensed data. Green areas show forest cover in 2000 and orange shows areas that experienced loss between 2000-2018. The black lines encompass the marine transgression zone for the purpose of this study, defined as 30 m landward from the edge of tidal marsh.

**Figure 2.**
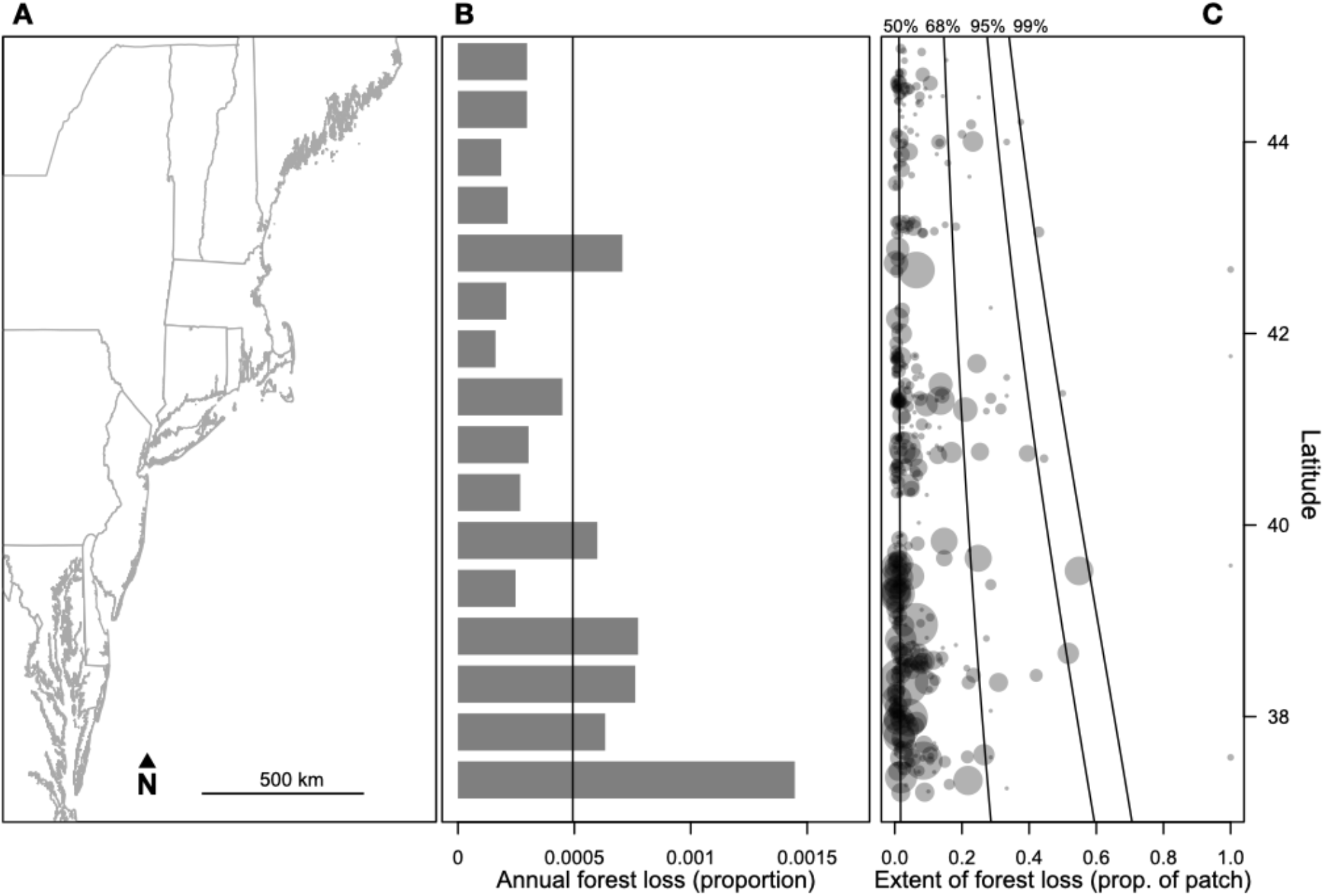
The creation of ghost forests across the Northeast U.S. between 2000-2018. **A** shows the state boundaries for the geographic extent of the study area. **B** shows the observed forest loss, as a proportion of the total area of forest within 30 m of tidal marsh, for 0.5-degree latitudinal bands. The black line shows the mean for annual forest loss across the study area. **C** shows the proportion of the marine transgression zone that experienced loss for all patches that experienced at least some loss (gray dots; larger dots represent larger patch areas). Black lines show the percentiles for the proportion of loss below which 50%, 68%, 95%, and 99% of the patches belong.

## Methods

I quantified forest loss in the marine transgression zone of the eastern U.S. between 2000-2018 using Hansen et al.’s (2013, 2018) Global Forest Change dataset (Figures 1, 2, S1). This region is the focus of research to better understand the responses of coastal ecosystems to sea-level rise (Lentz *et al.* 2016) and a recent review of research needs for understanding how marine transgression drives land conversion (Kirwan & Gedan 2019). It is also the focal region for the Atlantic Coast Joint Venture, a large-scale conservation initiative focused on coastal species and ecosystems (U.S. Fish and Wildlife Service 2019). I quantified the relative importance of a set of physical and ecological factors that have been hypothesized to drive marine transgression. I quantified values for these physical and ecological variables and extracted patch-scale trends in coastal forest loss using Google Earth Engine (Gorelick *et al.* 2017). I then quantified trends in forest loss and the strength of the potential predictive variables using Bayesian hierarchical models that incorporated approaches for protecting against overfitting and quantified parameters with full uncertainty bounds. I discuss this approach in more detail below.

### Hypothesized drivers of marine transgression

#### Forest cover and loss

I quantified forest cover and loss in the transgression zone using the Hansen Global Forest Change dataset v1.6 (Hansen *et al.* 2013, 2018). This dataset measures for each pixel (30-m resolution) whether there was evidence of forest loss between 2000-2018. I used this dataset to quantify for each patch (see <u>tidal marsh extent and transgression zone</u> below) the area of the transgression zone that experienced forest loss. For the analyses in this paper, I defined forest as greater than 20% tree cover to 1) distinguish loss of stands from loss of individual trees, and 2) be consistent with the National Land Cover Database, which I used to measure evergreen cover (see below).

#### Tidal marsh extent and transgression zone

I delineated tidal marsh patches according to Wiest et al. (2016), who used a 50-m buffering approach that groups marshes into biologically-relevant units using the U.S. Fish and Wildlife Service’s National Wetlands Inventory (https://www.fws.gov/wetlands/nwi/). I defined the transgression zone as the first 30 m from the landward boundary of each marsh patch to capture the frontier of marine transgression. I conducted analyses of drivers of forest loss at the patch-level, with variables summarized as the mean, sum, or proportion of the patch, as appropriate, using functions for dataset manipulation and image reduction (see code in **Supplementary code**; all spatial data was in projection EPSG:4269). Thirty meters from the landward boundary of tidal marsh is a useful distance for comparing across larger areas, as there is substantial variation in the extent of marine transgression. I also tested sensitivity to the choice of this distance by comparing the results using 30 m to the results using 100 m. While 30 m is a useful choice because it captures forest loss well across the study area, 100 m might be more appropriate for areas, like the southern latitudes of the study area, that have experienced greater loss. The choice of distance, however, did not alter inferences about the effect or importance of predictive variables (Figure S2), or the overall rarity of forest loss in the transgression zone (0.88% for 30 m vs. 1.5% for 100 m), and all results reported are for 30 m.

#### Upland slope

The slope of the upland immediately adjacent to tidal marshes has been shown by numerical models (Kirwan *et al.* 2016) and field studies (Anisfeld *et al.* 2017) to be a significant determinant of the extent and rate of marine transgression. I, therefore, expected shallower slopes to be correlated with more forest loss. I quantified the mean slope of each patch’s marine transgression zone using the U.S. Geological Survey National Elevation Dataset (1/3 arc-second resolution). This dataset was also used in a recent analysis of the dynamic responses of coastal ecosystems over the same study region (Lentz *et al.* 2016). I calculated slope from this dataset in Google Earth Engine (function ‘ee.Terrain.slope’; see code in **Supplementary code**).

#### Proportion of evergreen cover

The community composition of coastal forest varies by latitude, with coniferous species being dominant in the northern and southern areas of the study region. Previous research has suggested that forest loss from marine transgression could largely be a result of lack of recruitment once older individuals die (Kirwan *et al.* 2007), and more generally, that forest persistence can be driven by re-sprouting (Bond & Midgley 2001), a trait that woody conifers rarely possess (Clarke *et al.* 2013). I therefore expected more forest loss in patches dominated by conifers. I quantified the proportion of the forest in each patch’s marine transgression zone that was evergreen using the U.S. Geological Survey’s National Land Cover Database (NLCD; Yang *et al.* 2018; 30-m resolution). For each patch, I quantified the proportion of pixels that were classified as either 42 (evergreen) or 43 (mixed).

#### Rate of sea-level rise

Sea-level rise is the primary driver of marine transgression and has been associated with several indicators of the conversion of coastal forest to wetland (Desantis *et al.* 2007; Doyle *et al.* 2010; Smith 2013). Regional variation in the rate of sea-level rise, however, has never been investigated as a potential driver of trends in marine transgression. I specified the rate of sea-level rise (mm/year) for each patch according to the closest tide gauge from the National Oceanic and Atmospheric Administration (NOAA; https://tidesandcurrents.noaa.gov).

#### Peak storm surge during Hurricane Sandy

Previous research on the responses of coastal forest to sea-level rise suggested that loss will likely occur stepwise in response to storm-induced mortality (Brinson *et al.* 1995; Kirwan *et al.* 2007). I was able to test this prediction as an extreme storm, 2012’s Hurricane Sandy, affected the center of the study area during the study period. I determined the peak storm surge during Sandy for each patch according to the closest gauge from the U.S. Geological Survey’s (USGS) survey of water levels during the storm (McCallum *et al.* 2013). Each measurement represents the peak storm tide experienced between October 29 and 30, 2012, in ft. above the North American Vertical Datum of 1988. The USGS deployed gauges at 147 locations across the study region. Patches that occur north or south of the geographic range of these sensors were assigned the value from the northern-most (43.544722, North American Datum of 1983) or southern-most gauge (36.700430), as appropriate.

#### Patch area

Brinson et al. (1995) suggested several mechanisms by which coastal forest could maintain positive feedbacks that would prevent or slow the transition from forest to wetland, such as the maintenance of fresh groundwater. These feedbacks should be stronger for patches that have more trees, leading to the expectation of less forest loss in larger patches. I quantified patch area as the as the sum of pixels in the marine transgression zone that was covered by forest in 2000 (see <u>Forest cover and loss</u> above).

#### Geomorphic setting

Tidal marshes in the eastern U.S. vary by important biophysical processes such as hydrodynamics, sediment supply, and ecological communities (Cahoon *et al.* 2009; Wiest *et al.* 2016), which are often grouped into geomorphic settings (Reed *et al.* 2008). I used the categories from Wiest et al. (2016), which are based on Reed et al. (2008) and include open coast, back-barrier lagoon, estuarine embayment, estuarine brackish marsh, tidal fresh marsh, and nontidal brackish marsh. I did not have *a priori* expectations for in what directions each geomorphic setting would influence forest loss, given the complexity of the relevant hydrodynamics and ecological processes. I instead included geomorphic setting as a categorical variable to account for any differences between settings that are not captured by the other variables in the model.

### Statistical analyses

#### Quantifying variables that predict forest loss

I quantified the strength of variables that potentially explain patterns of coastal forest loss using a Bayesian statistical model in a generalized linear modeling framework. I developed this model using continuous model expansion guided by posterior checks (Gelman *et al.* 2013) and a simulation model to test key aspects of the model design and ensure parameter identifiability. Histograms of the proportion of each coastal forest patch that experienced loss showed a strong peak at zero, suggesting that a zero-inflation model was warranted to appropriately model rare events. I therefore specified the model in two parts: one that aims to explain variation in the probability of at least some forest loss (0 or 1), and one explains the proportion of forest loss, at the patch-level, given that there was some loss. My approach was a zero-additive, rather than zero-inflated, because I only used patches that had at least some loss when quantifying the proportion of loss (Martin *et al.* 2005; see model code in **Supplementary code**). The benefits of this approach include 1) running the Monte-Carlo-Markov-Chain (MCMC) estimation on smaller graph sizes, resulting in realistic run times; 2) an intuitive interpretation in which one model quantifies how the set of hypothesized drivers explains the probability of any forest loss, and the other quantifies how this set of drivers explains the proportion of loss – i.e. the size of the ghost forest, given that one was created; and 3) ensuring that the distribution of the dependent variable, forest loss, was appropriately explained by the sampling distributions of the models. I used a simulation based on known values to show that the model specification was able to recover unbiased estimates of regression coefficients and make unbiased predictions of forest loss (Figures S3, S4, S5; see *Simulation model* below).

I first modeled the probability of at least some forest loss using a Binomial distribution as follows:

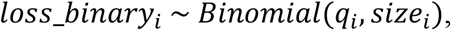

where *loss_binary_i_* is an index of whether any pixels in patch *i* experienced loss (1 is no loss; 0 is at least some loss), *q* is the probability that a given pixel in a patch does not experience loss, and *size* is the number of pixels in a forest patch. I related *q* to *π*, the probability that a given pixel in a patch experiences loss, by the following relationship:

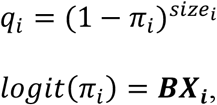

where ***BX_i_*** is the coefficient matrix of the regression equation. Modeling the probability of a patch *not* experiencing loss in this way made it possible to treat loss as a binary variable while also taking into account the size of the patch, thereby accounting for the fact that large patches have more opportunities for loss.

I modeled the proportion of the marine transgression zone that experienced forest loss, given that there was as least some loss, using a Binomial distribution with normally distributed overdispersion:

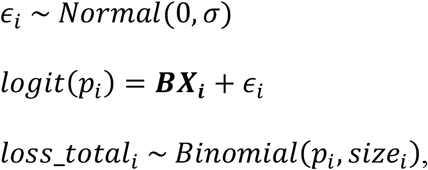

where *p*_*i*_ is the probability that a given pixel in patch *i* experiences loss, given that there was at least some loss in the patch, *ε* is variation in the probability of loss that is not explained by the sampling variation of the Binomial distribution, and *σ* is the variance of unexplained variation. Because a logistic link was needed to incorporate regression coefficients (see model code in **Supplementary code**), the resulting overdispersion was a logistic-normal distribution, which quantile-quantile plots showed was an appropriate description of the residuals.

Both model structures described above explicitly account for the fact that larger patches have more opportunities to experience loss. Simulations demonstrated that this model specification does successfully account for the effect of area on binomial probabilities while also allowing area to potentially affect loss through a biological mechanism (e.g. see <u>Patch area</u> above; Figure S3).

Once I established the basic error structures for the models, I allowed both the probability of some loss, *π*, and the proportion of loss at the patch-level, *p*, to vary according to the variables described above, corresponding to the hypotheses for which factors drive forest loss. All of the variables considered had strong *a priori* justifications or prior evidence that they are likely to influence forest loss (see *Hypothesized drivers of marine transgression* above). Accordingly, my approach for avoiding overfitting was to use Bayesian variable selection (O’Hara & Sillanpää 2009), which specifies a prior distribution that shrinks estimates toward zero, representing the prior belief that most variables are likely to have a moderate to small, but non-zero, effect. This adaptive shrinkage approach (*sensu* O’Hara & Sillanpää 2009) is similar to “spike and slab” approaches, but in this case a spike is not specified as there were strong prior beliefs that all variables were likely to influence loss. Instead, I specified a t distribution with 4 degrees of freedom, which shrinks estimates toward zero while allowing some variables to potentially have a large effect. I tested the sensitivity of the parameter estimates to this variable selection approach by also fitting two alternative models: one that used a normal distribution instead of a t distribution, and one that treated variables as fixed effects, both of which gave similar results.

Posterior checks showed that a quadratic term for area was warranted, as loss increased sharply for very small patch sizes (Figure S6). I also allowed for two interaction effects: between sea-level rise rate and slope, and between the proportion of evergreen cover and latitude, to represent how I expected these variables to influence loss. I expected sea-level rise to have a larger influence at shallower slopes, with potentially little to no effect for the steepest slopes. I allowed the effect of the proportion of evergreen cover to vary by latitude to account for the fact that the conifer forests in the northern and southern parts of the study area have different species composition (dominated by lobblolly pine [*Pinus taeda*] in the south vs. eastern white pine in the north [*Pinus strobus*]; (Thompson *et al.* 1999).

Posterior estimates of the normal random effects, which accounted for overdispersion relative to the binomial model, suggested that variance decreased with slope and increased with area. I therefore allowed the variance parameter for the proportion of loss to vary by slope and patch area (see model code in **Supplementary code**). The final model for the proportion of loss, given at least some, included 1) a binomial distribution and normal random effect that modeled the proportion of pixels in each patch that experienced loss, 2) regression coefficients for variables that relate to the factors hypothesized to drive forest loss, and 3) an equation that models the variance of loss by slope and area.

I compared the relative influence of the physical and ecological variables on forest loss by standardizing each variable on the same scale (subtracting the mean and dividing by 2 standard deviations; Gelman 2008) and comparing the magnitude of their posterior estimates. I quantified the importance of including these potential drivers of marine transgression in the models, as a group, by comparing the patch-level posterior predictions of models with and without drivers to the observed rates of loss (Figure S7). For the model of the probability of at least some loss, a binary variable, I plotted the empirical Receiver Operating Characteristic (ROC) curve, which quantifies the trade-off between specificity and sensitivity for binary predictions (Figure S8). I also calculated the area under the ROC curve (AUC), which provides a metric that compares the performance of a model with drivers to the performance of a model that has no predictive ability (AUC = 0.5). To obtain measures of the model’s predictive ability that are more informative than a single value (Pontius & Parentier 2014), I examined the shape of the ROC curve to check for systematic patterns of underperformance and calculated the confidence bounds of AUC by propagating the uncertainty of the model’s posterior predictions. Semi-variograms of the model residuals by distance did not show spatial patterns after variables that potentially influence loss were accounted for.

#### Quantifying recent trends in rates of forest loss

I quantified potential trends in the rate of forest loss across latitudinal bands (0.5 decimal degrees). A positive latitudinal trend in the total area of loss per year would signal accelerating loss. I specified a hierarchical model that estimated band-specific trends, which were related to each other by a global distribution with a mean trend for the study area and a parameter that describes the variance in trends among bands (see model code in Supplementary code). The model allowed each band to have an independent parameter for sampling variation. I also used this model structure to estimate potential trends in the number of patches in each year that experienced loss, which gives a measure of the number of discrete loss events over time.

I conducted all statistical analyses in JAGS (Plummer 2017) in R v4.0.3 (R Development Core Team 2021) using the R2jags package (Su & Yajima 2015). I used uninformative priors for all parameters except for the Bayesian variable selection described above. I checked that the parameter estimates were not sensitive to the specific choice of priors by fitting models using more and less informative prior information (see model code in **Supplementary code**). I ran the model for 100,000 iterations after an 100,000-iteration burn-in, with no thinning. To check convergence, I ran three MCMC chains and checked that the R-hat values for each parameter were < 1.01 (Brooks & Gelman 1998).

### Simulation model

I specified a simulation that generated known values with the key features of the empirical data on forest loss (see simulation code in Supplementary code): I generated data from 5000 patches, less than the 7350 patches of the observed dataset, to ensure that the model performed adequately with fewer data. I specified zero inflation in line with the observed data (~20% of patches experienced at least some loss). I used the simulation primarily to ensure that three aspects of the model worked as expected: 1) that predictions of binomial probabilities of loss were unbiased (Figure S4), 2) that the model was able to disentangle area effects that were a result of potential biological or physical mechanisms from the fact that larger patches have more opportunities to experience loss (Figure S3), and 3) that the model could recover unbiased parameter estimates for a variable that is correlated with area (Figure S5). For this latter check, I introduced a slight (~10%) correlation between patch area and another variable, as observed in the empirical data (e.g. for latitude; see Table S1).

## Results

### Spatio-temporal trends in rates of forest loss

Between 2000-2018, the creation of new ghost forests was a relatively rare phenomenon across the northeastern U.S., with greater than 80% of the observed forest loss concentrated in less than 0.84% of marshes (62 marsh patches) and 95% of loss concentrated in less than 3.86% of marshes (284 patches; Figure 3). During this time period, 0.88% of the entire forested area of the transgression zone experienced loss (1000 ha; Figures 1, S1). Regionally, the smallest area of loss was 0.29% of the forested transgression zone (0.26 ha) in northern latitudes and the largest area was 2.57% at lower latitudes (16.81 ha; Figures 2, S1). There were few examples of patches that experienced loss across more than 80% of their forested area. Across most of the latitudinal gradient of the study area, greater than 95% of patches experienced less than 50% loss (Figure 1). While much less common than loss, the transgression zone also experienced forest gain, with a peak of 0.49% at lower latitudes (Figure S1).

**Figure 3.**
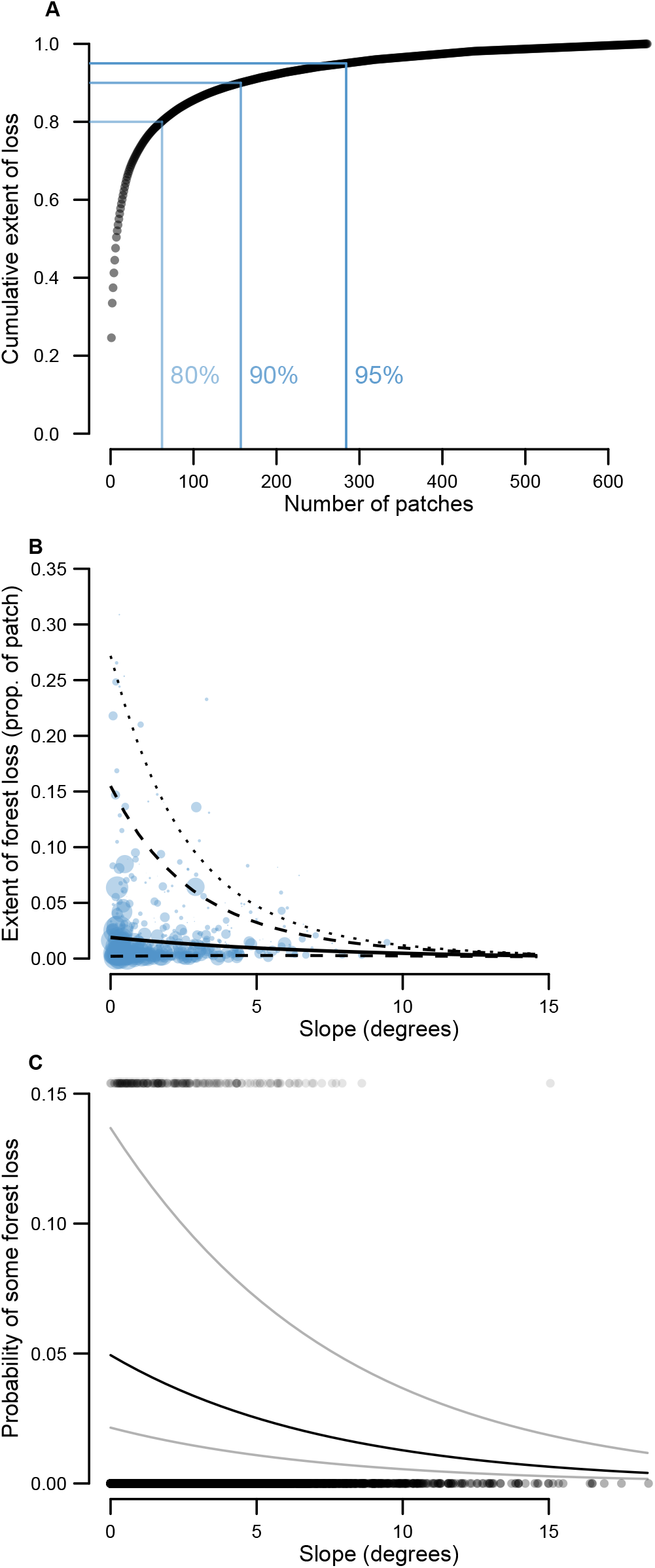
The distribution and likelihood of the creation of ghost forests. Plot **A** shows the cumulative extent of coastal forest loss by the total number of patches, with patches ordered from most to least loss. Grey dots show the cumulative extent of loss as each patch is added, as a proportion of the total loss observed in the dataset. Blue guides show the 80%, 90%, and 95% percentiles of the total loss observed. Only the first 600 patches are shown. Greater than 80% of the loss is concentrated in less than 100 patches, and 95% of the total loss is clustered in less than 300 patches. Plot **B** shows the model predictions for the proportion of forest within the marine transgression zone that experienced loss across the range of values for slope. The solid black line shows the mean proportion of loss for median-sized patches. Dashed lines show the 95% bounds and the dotted line shows the upper 99% bound. Blue dots show the observed proportions of loss for patches that experienced loss (larger dots represent larger patch areas). Both the mean and variance of the proportion of loss increase for shallower slopes, with a particularly steep increase in the variance. Plot **C** shows the probability of forest loss in the marine transgression zone as a function of slope, with all other variables except patch area held constant at their means. The solid black line shows the relationship between loss and slope for median-sized patches, and the gray lines show the relationship for the lower 0.25 and upper 0.75 quantiles of patch area. Gray dots show the observed loss for each patch (0 for no loss or 1 for some loss).

The mean probability and extent of forest loss between 2000-2018 increased with shallower slopes, but this increase, while statistically detectable, was small (Figure 3b, c). In contrast, the variation in the extent of forest loss increased sharply with shallower slopes (Figure 3b). This pattern suggests that the observed trend of greater loss at lower latitudes is driven by increases in the variance rather than the mean, as slope tends to increase with latitude across the study area (Pearson’s correlation coefficient: 0.50; 95% confidence interval: 0.48, 0.52).

I found little evidence of acceleration in the rate of forest loss between 2000-2018. The posterior probability that annual forest loss is accelerating across the study area was 0.19, and all estimates of loss by latitudinal bands overlapped zero (Figure 4a). No latitudinal bands were outliers from the overall trend in the rate of forest loss for the study area (Figure 4a). While the current weight of the evidence suggests a slowing of the rate of loss (Figure 4a), the 95% bounds for the overall trend overlapped zero (−0.24 ha [95% credible interval: −0.82, 0.38]). The trend in the number of discrete loss events per year was negative, suggesting a tendency toward fewer new ghost forests between 2000-2018 (parameter estimate for annual trend in loss events per year: −0.07% [−0.12, −0.02]). None of the latitudinal bands show trends that differed substantially from this overall trend of fewer new ghost forests, with several latitudinal bands also showing statistically detectable deceleration in the number of loss events (Figure 4b).

**Figure 4.**
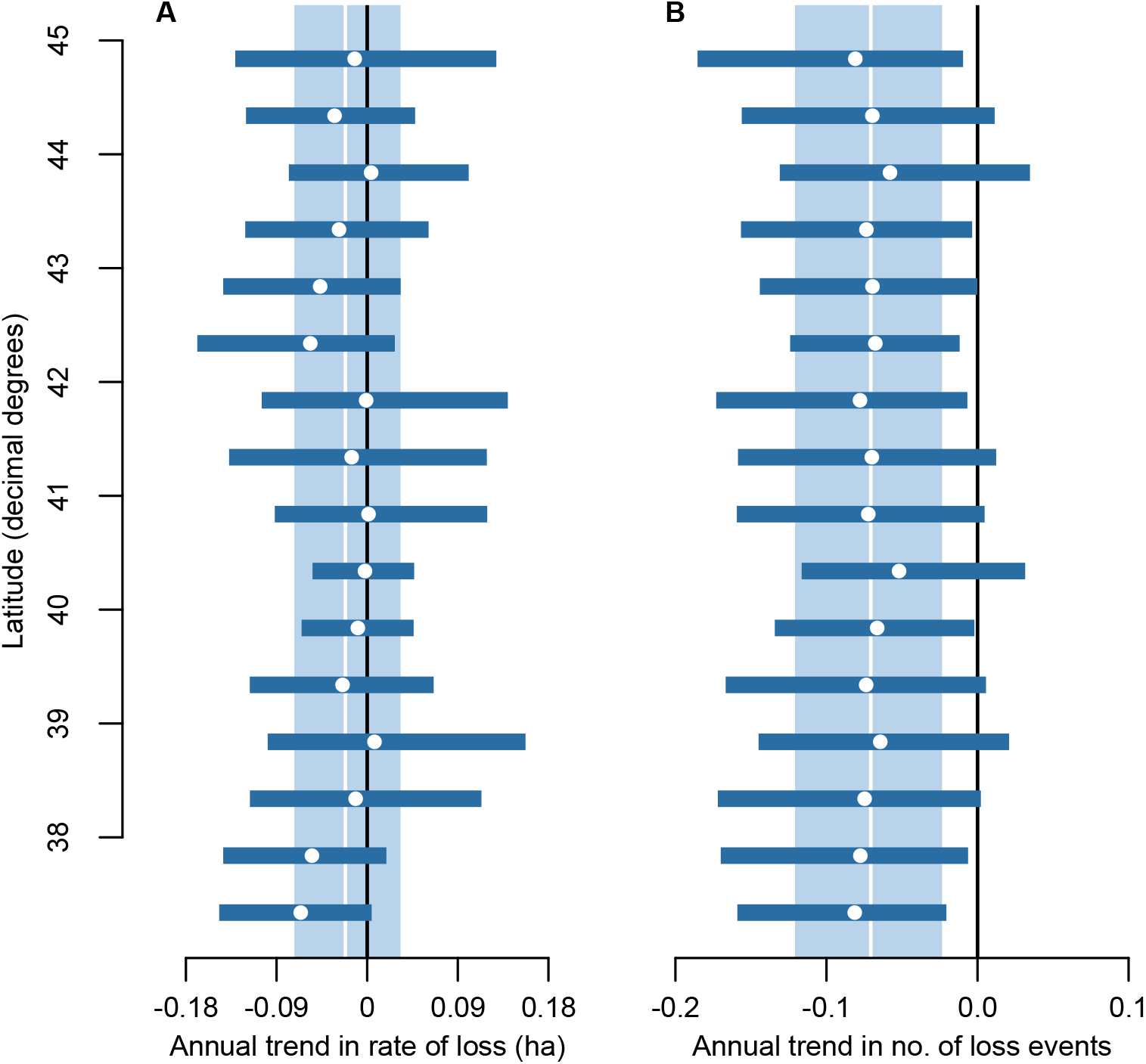
Trends in the rate and frequency of the creation of ghost forests between 2000-2018. Plot **A** shows parameter estimates for the trends in the annual rate of loss in the marine transgression zone for 0.5-degree latitudinal bands across the study area. Plot **B** shows parameter estimates for the trend in the number of loss events per year. For both plots, white dots show the posterior mean for the trends for each band and blue bars show the 95% credible intervals. The vertical white lines show the overall trend for the study area and the light blue shaded areas show the corresponding 95% credible intervals. For reference, a vertical line at zero shows the expectation for no trend in the creation of ghost forests. The 95% bounds for the trend in the number of forest loss events across the study area are less than zero, suggesting that the number of loss events per year declined over the study period.

### Factors that influence forest loss

Physical, ecological, geomorphic, and disturbance variables all explained observed patterns of forest loss: for all variables, the 95% credible intervals did not include zero for either the presence (i.e. probability) or extent of loss (Figure 5) once interactions were accounted for (Figure 6). All variables and their interactions influenced loss in the direction expected by *a priori* predictions (Figures 5, 6). As expected, the effect of the rate of sea-level rise on forest loss was moderated by slope, with a greater positive effect at shallower slopes (Figures 5, 6). I also found evidence for an interaction between latitude and the extent of conifer cover, which was correlated with both a greater probability and extent of loss (Figures 5, 6c). In both cases, the effect of evergreen cover was most positive (i.e. greater probability or extent of loss) at lower latitudes (Figures 5, 6c).

**Figure 5.**
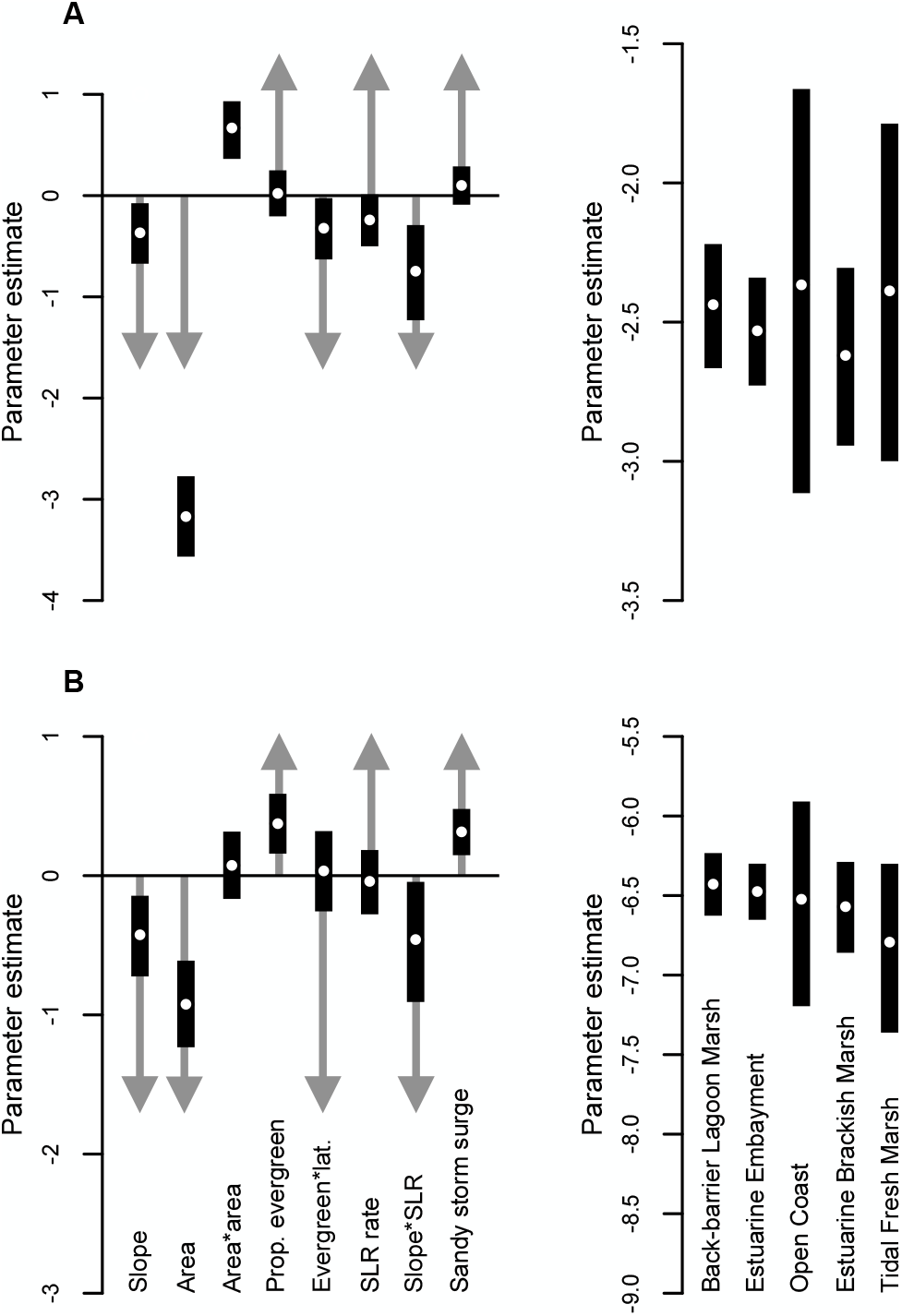
Parameter estimates of the coefficients for factors that potentially explain the likelihood and extent of ghost forests. Plots show parameter estimates for variables hypothesized to influence: **A** the proportion of the marine transgression zone that experienced loss, given at least some, and **B** the probability of at least some forest loss. White dots show posterior means and black bars show the 95% credible intervals. Gray arrows show the *a priori* predictions for the direction of the influence, before interactions are accounted for. All variables for which the credible intervals do not overlap zero differed in the expected directions, and the bulk of the probability density for all variables is in the expected direction, once the influence of interaction effects is taken into account (Figure 6 better shows the influence of these interaction effects than the separate coefficient estimates shown here). A prediction is not shown for the interaction effect for area because this effect was found to improve the model during posterior checks and therefore I did not make an *a priori* prediction.

**Figure 6.**
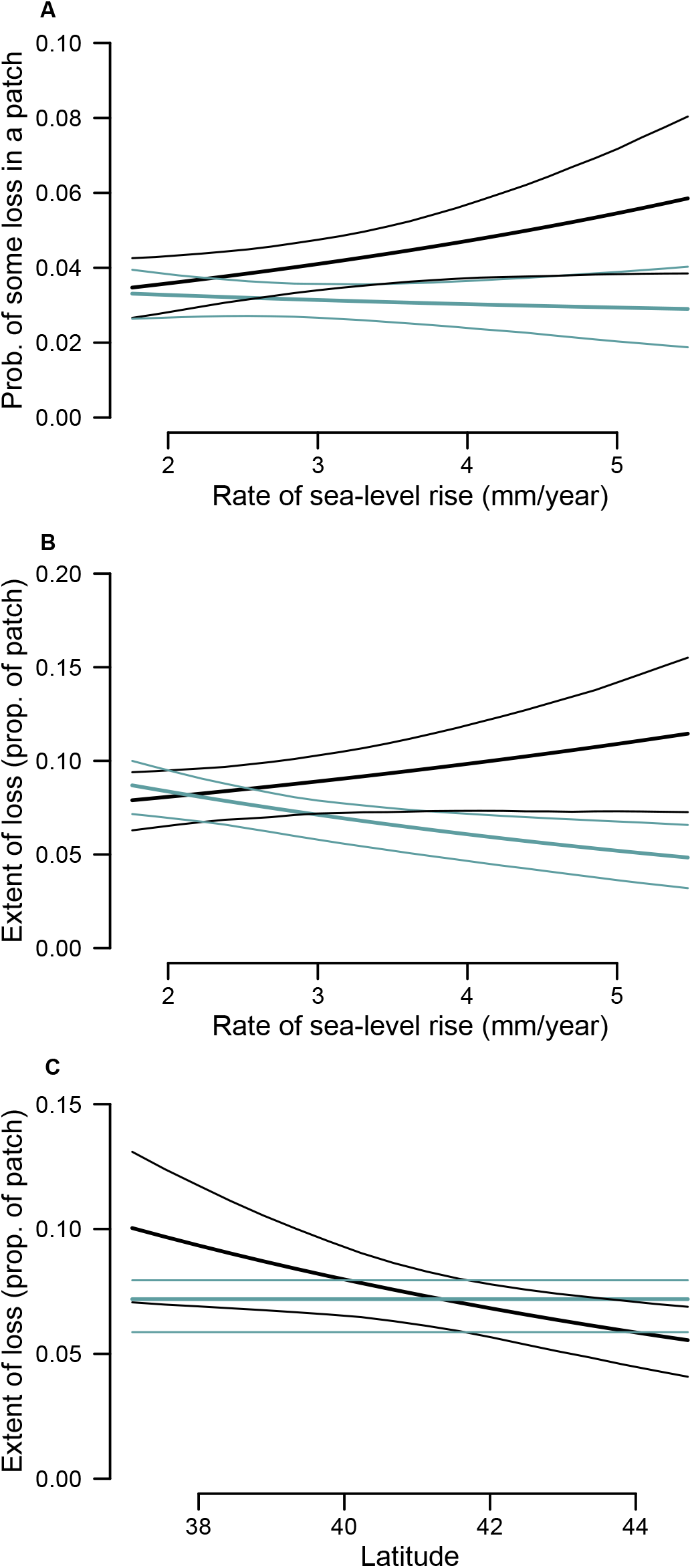
Predicted relationships for key interaction effects in models of the probability (**A**) and extent (**B** and **C**) of ghost forest formation. **A** shows the how relationship between the rate of sea-level rise and the probability of forest loss in the marine transgression zone is affected by slope. Blue lines show the posterior mean (thicker line) and 95% credible intervals (thinner lines) for patches with the mean observed slope and black lines show this measure for patches with the minimum observed slope (black lines). Plot **B** shows the how relationship between the rate of sea-level rise and the proportion of forest loss in the marine transgression zone, given at least some, is affected by slope. Blue lines show the posterior mean (thicker line) and 95% credible intervals (thinner lines) for patches with the mean observed slope and black lines show this measure for patches with the minimum observed slope (black lines). Plot **C** shows the how relationship between the proportion of conifer cover and the proportion of forest loss, given at least some, is affected by latitude. Blue lines show the posterior mean (thicker line) and 95% credible intervals (thinner lines) for patches with no conifer cover and black lines show this measure for patches with complete conifer cover (black lines).

The height of peak storm surge during Hurricane Sandy was one of the strongest predictors of the probability of a patch experiencing some loss (Figure 5b), with higher surge predicting a greater probability of loss. Surge height during Sandy was not correlated with a greater extent of loss in patches that experienced at least some, however (Figure 5a), suggesting that storm surge caused forest loss but is not the primary driver of extensive ghost forests. The total area of forest in a patch was a strong predictor of both the probability and extent of loss (Figure 5), with patches containing less total forest experiencing greater loss (Figures 5a, S2) after accounting for the fact that large patches have more opportunities for loss (Figure S3).

Coastal areas that are classified as back-barrier lagoon or estuarine embayment geomorphic settings experienced the greatest forest loss (0.99% and 0.98% respectively), with estuarine brackish marsh experiencing the least (0.41%; Figure S9). I also found evidence that these differences in geomorphic setting remained after accounting for the other biophysical variables: for both the probability and extent of forest loss, the standard deviation of the effects of geomorphic setting were well above zero with all other variables in the models (Figure S9).

The models of the probability and extent of forest loss both performed better when independent variables for the physical, ecological, geomorphic, and disturbance variables were included (Figures S7, S8), providing further evidence that these variables are important for understanding and predicting marine transgression. For the model of the probability of forest loss that included all variables, the area under the Receiver Operating Characteristics curve (AUC), was 0.737282 (95% confidence interval: 0.69, 0.78). Compared to an AUC value of 0.5 for a model with no predictive ability, this value suggests that including biophysical variables improves the amount of information with which to make predictions. The model’s improvement over no predictive ability was consistent over the range of forest loss probabilities (Figure S8). For the model of extent of loss, the posterior predictions from a version of the model that included biophysical variables performed substantially better than a model without variables, especially for areas more loss (Figure S7).

## Discussion

Ghost forests are one of the most easily observable, and memorable, indicators of marine transgression, but systematic analysis across the entire northeastern U.S. revealed they are a relatively rare phenomenon. My analysis also found a clear trend of greater forest loss at lower latitudes, a pattern that has been previously hypothesized or noted anecdotally (e.g. Kirwan & Gedan 2019). This study is the first to provide specific evidence for potential drivers of this trend. One potential driver is that lower latitudes have a greater extent of areas with shallower slopes, which results in more opportunities for rare events, like ghost forests, to occur. In this case, slope is a driver of observed patterns of loss, but slope, *per se*, is not a strong predictor of whether an area is likely to have experienced recent loss. As marine transgression progresses, however, it is likely that slope will become a stronger predictor of forest dieback (Kirwan *et al.* 2016) – i.e. slope will be predictive of the mean extent of forest loss – but the results reported here suggest that this point has not yet been reached.

The other potential driver of observed latitudinal trends is that lower latitudes are dominated by conifers (Thompson *et al.* 1999), which were associated with a greater probability and extent of loss. The effect of conifer cover was also stronger at lower latitudes, which could be the result of differences in species composition and diversity or interactions with physical or hydrological factors. More research is needed to disentangle the potential physiological, hydrological, or ecological drivers of the apparent difference in resilience to sea-level rise between areas dominated by deciduous vs. coniferous species. Understanding the resilience of forest in general has profound implications for the timescales of marine transgression, especially when compared to coastal wetlands, which appear to be experiencing more rapid change (Field *et al.* 2016). Increasing evidence suggests mechanisms that would make coastal forest resistant to change in the short term (e.g. Fagherazzi *et al.* 2019; Kearney *et al.* 2019), which could cause temporary or long-term bottlenecks in the extent of native coastal wetlands, especially if persistent coastal forest gives a competitive advantage to non-native vegetation (Shaw *et al.* 2021).

With any remote sensing study there is a trade-off between making inferences across large scales and incorporating all of the factors that are potentially influential. Measures of forest loss, in particular, are reasonable targets for remote sensing and supported by a strong body of evidence (Hansen *et al.* 2013, 2018), but there are drivers that might be important that are not as easily obtained over large scales. While I was able to obtain a wide range of different data types here, there are several additional factors that I could not include that are known to influence some aspect of marine transgression. For example, measures of ground water discharge are likely important for understanding tree death and potential tipping points that govern dieback at the stand-scale (Brinson *et al.* 1995), and the seedbank can play a large role in determining the potential for regeneration of upland vegetation (Kottler & Gedan 2020). Studies that further elucidate these processes are essential for developing and testing the conceptual model of marine transgression (e.g. Brinson *et al.* 1995), while correlational studies like this one can provide broader regional context, including measures of relative importance among the large set of potential drivers.

The patterns illuminated by this study provide context for the observation that studies from higher latitudes found less evidence for marine transgression (e.g. Field *et al.* 2016) than studies from lower latitudes, some of which found extensive forest loss (e.g. Smith 2013). The simplest hypothesis for these regional differences is that forest loss is driven by shallower slopes at lower latitudes (Kirwan & Gedan 2019). My analyses showed greater support for alternative hypotheses, however, including that loss is driven by opportunities for rare events and the extent of conifer cover. The variables included in my analyses are likely appropriate proxies for the processes being quantified in more intensive field studies, which is supported by the result that all of the factors influenced forest loss in the expected direction, once interactions were accounted for. This results also highlights the importance of thinking broadly as well as deeply when developing studies or models of the drivers of marine transgression.

If remotely sensed measures of forest loss are biased for coastal areas, it could potentially bias analyses like those reported here. While no evidence for this bias currently exists, one possibility is that the pattern of dieback from saltwater inundation, which can occur in linear strips as opposed to patches, is biased high or low by the methods in Hansen (2013, 2018). If such a bias did exist, however, it would likely be consistent across the study area and therefore not have a large effect on correlational analyses. It is also unlikely that all observed forest loss is from saltwater inundation, as edge effects that are not specific to marine environments, though often overlooked in studies of ghost forests, likely contribute to the observed extent of dieback. If this is the case, ghost forests associated with marine transgression might be even rarer than the estimates here indicate.

The relative rarity of ghost forests, the relatively slow rate of forest loss, and the lack of evidence for acceleration cast doubt on whether marine transgression will progress rapidly enough to prevent the loss of tidal marsh specialists. The tidal marshes of the northeast U.S. are a hotspot of endemism for mammals and birds (Greenberg et al. 2006) such as the saltmarsh sparrow (*Ammospiza caudactus*), which is currently facing extinction by mid-century (Field *et al.* 2017b); Endangered on the International Union for the Conservation of Nature RedList). The current rate of conversion from uplands to tidal marsh is not likely sufficient to prevent extinction of saltmarsh sparrows and other specialists unless there are non-linearities or tipping points in the response of coastal forest to sea-level rise. The importance of these non-linearities or tipping points for understanding and predicting the extent of forest loss has been demonstrated by conceptual work (Brinson *et al.* 1995), modeling (Kirwan *et al.* 2007), and paleo-ecological evidence (Thomas & Varekamp 1991). The results here also suggest that the fingerprints of pulse events, like Hurricane Sandy, can already be found in recent trends at large scales. Despite these lines of evidence, there remains substantial uncertainty about how these pulse events will determine the timing and magnitude of the marine transgression and shoreline encroachment.

Projection modeling is one powerful and commonly used tool for better understanding how press and pulse factors will determine the extent of marine transgression, forest dieback, and wetland migration across time and space (e.g. Kirwan *et al.* 2007; Wu *et al.* 2015). The correlational analyses here highlight how factors that are not typically included in these efforts can be the strongest predictors of forest loss, suggesting that projections could be improved by integrating a wider range of ecological and disturbance factors, especially forest composition and the effects of large storms. Previous research has also demonstrated the importance of incorporating feedbacks between human communities and physical and ecological factors (Field *et al.* 2017a; Kirwan & Gedan 2019). A key question is whether, when all of these factors are considered, rates of marine transgression will be sufficient for the persistence of the unique plant and animal communities of higher elevation tidal marsh (Field *et al.* 2016). The plant and animal species of these “high marsh” areas are potentially valued highly by coastal communities and can serve as a central component of engagement and awareness efforts that aim to protect coastal ecosystems (Field *et al.* 2017a). Tidal marshes, as we know them, face an uncertain future in the northeast U.S. and globally over the next century of accelerating sea-level rise (Gedan *et al.* 2011; Kirwan *et al.* 2016). Widespread evidence for recent changes to these ecosystems (e.g. Donnelly & Bertness 2001; Field *et al.* 2016) highlights the urgency of interdisciplinary and synthetic analyses and models that can advance understanding of marine transgression and its role in determining the future of coastal ecosystems.

## Acknowledgements

This work was funded by the National Socio-Environmental Synthesis Center (SESYNC), under funding received from the National Science Foundation (DBI-1052875a).

## Notes

### Competing Interest Statement

The authors have declared no competing interest.

