## Supplemental figures and code for "The creation of ghost forests is driven by physical, ecological, and disturbance factors but remains a rare phenomenon"

### Supplementary tables and figures

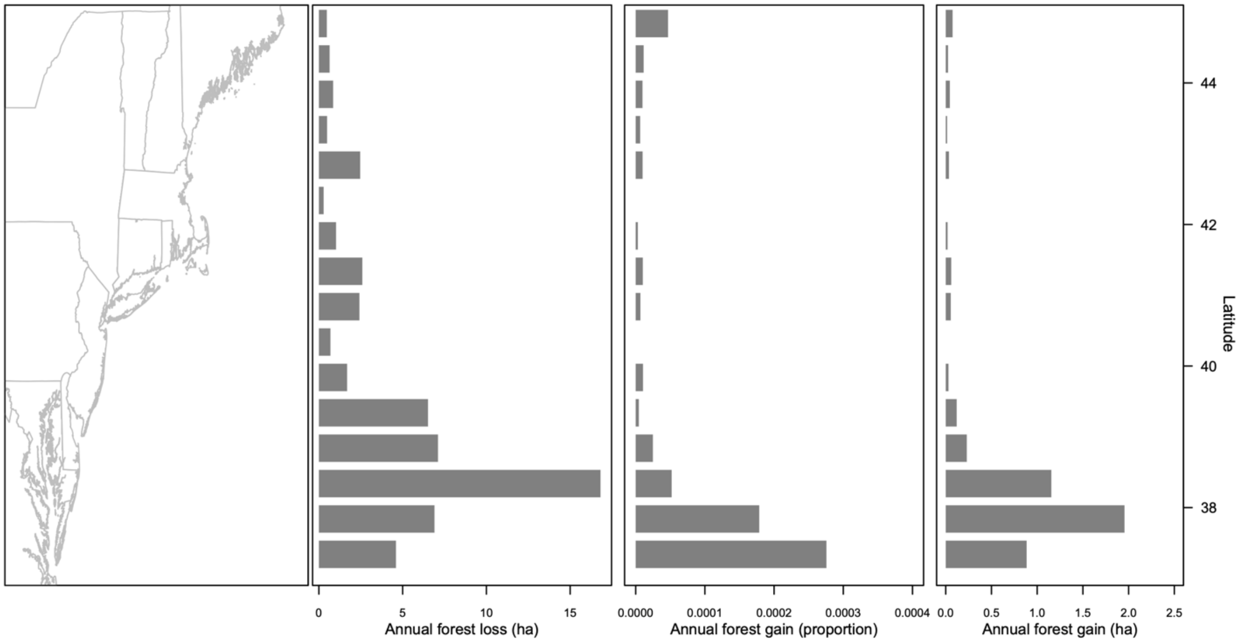


Figure S1. Observed forest loss across the Northeast U.S. between 2000-2018. The left panel shows the state boundaries for the geographic extent of the study area. The two middle and right panels show measures of observed forest change within 30 m of tidal marsh for 0.5-degree latitudinal bands. Measures shown are area lost per year (in ha), area gained per year (as a proportion of the total area of forest within 30 m), and area gained per year (in ha).

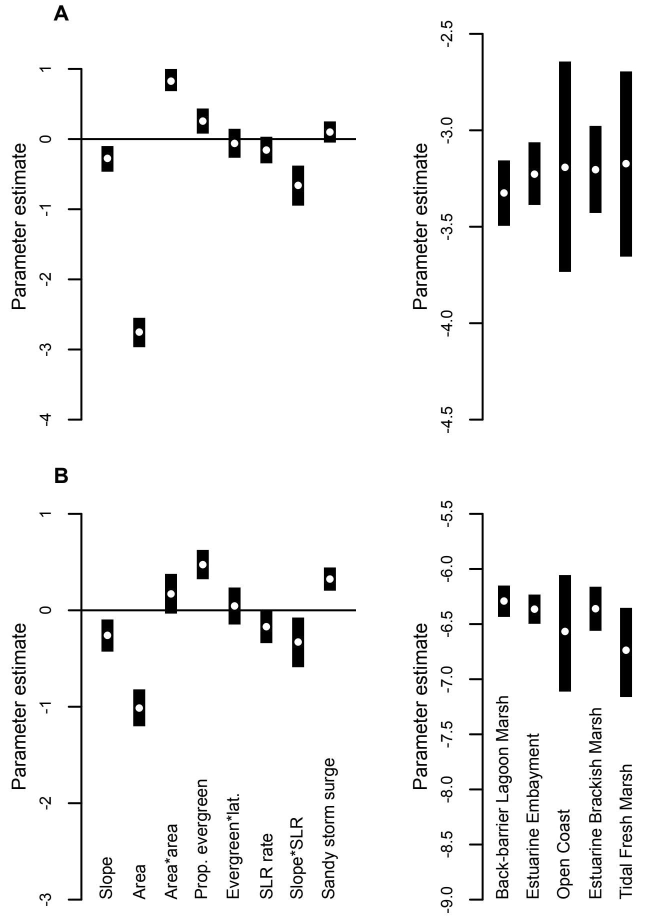


Figure S2. Parameter estimates of the coefficients for variables that potentially explain the likelihood and extent of the formation of ghost forests, using a 100 m definition for the extent of the marine transgression zone instead of the 30 m distance used for the primary results. Plots show parameter estimates for variables hypothesized to influence: **A** the proportion of the marine transgression zone that experienced loss, given at least some, and **B** the probability of at least some forest loss. White dots show posterior means and black bars show the 95% credible intervals.

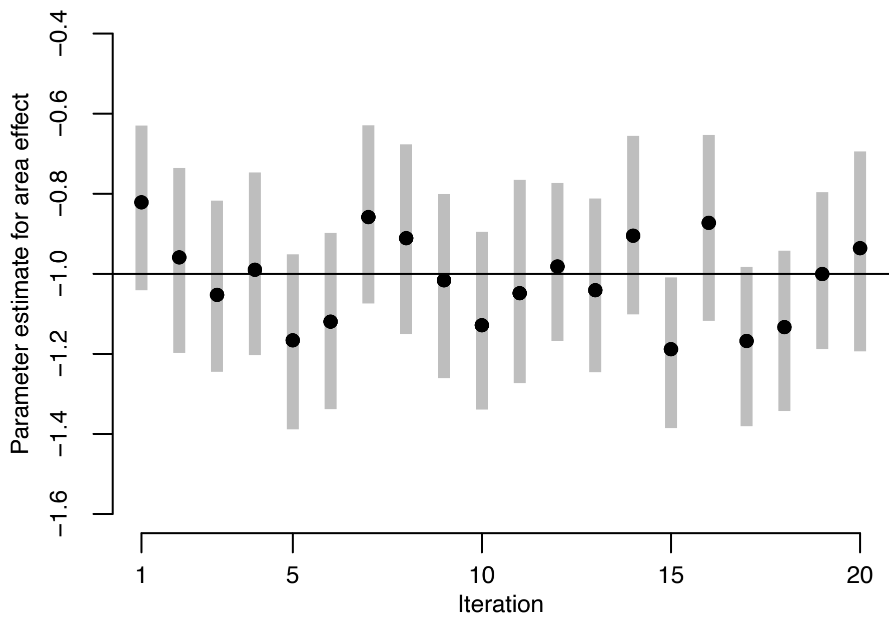


Figure S3. Parameter estimates for the effect of area on forest loss from a Bayesian hierarchical model of simulated data (see **Methods**). The value of the regression coefficient from the simulation that generated the data (-1) is shown as a horizontal black line. Black dots show the posterior means for 20 simulated datasets. Gray bars show the 95% credible intervals. As expected, estimates from simulated data were centered around zero, and approximately 95% of the credible intervals overlapped zero, demonstrating that the model was able to distinguish a true effect of area from the fact that larger forest patches had more opportunities for loss.

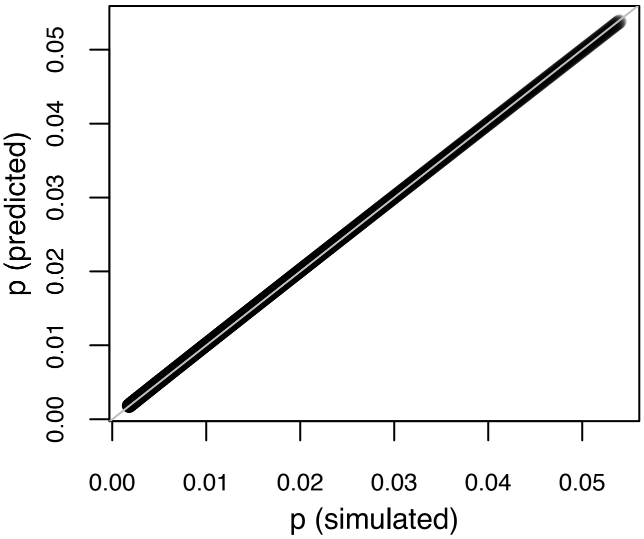


Figure S4. Comparison of the known values (simulated) for the probability of forest loss, *p*, and predicted values from a Bayesian hierarchical model (see **Methods**) that used the simulated data points. The relationship between predicted and simulated values falls along the 1:1 line (show in gray), suggesting that the model was able to recover unbiased estimates of the probability of forest loss using a simulated dataset with the same properties as the observed data for forest loss.

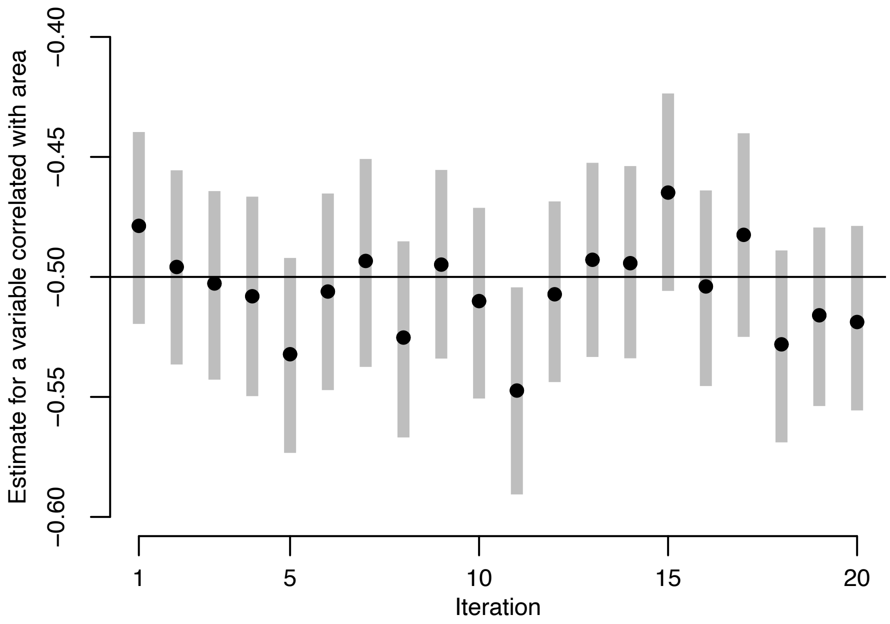


Figure S5. Parameter estimates for an effect that is correlated with area (r ~ 0.10) from a Bayesian hierarchical model of simulated data (see **Methods**). The value of the regression coefficient from the simulation that generated the data (-0.5) is shown as a horizontal black line. Black dots show the posterior means for 20 simulated datasets (true value is 0.20). Gray bars show the 95% credible intervals. As expected, estimates from simulated data were centered around zero, and approximately 95% of the credible intervals overlapped zero, demonstrating that the model structure can recover parameter estimates for variables that are potentially correlated with area.

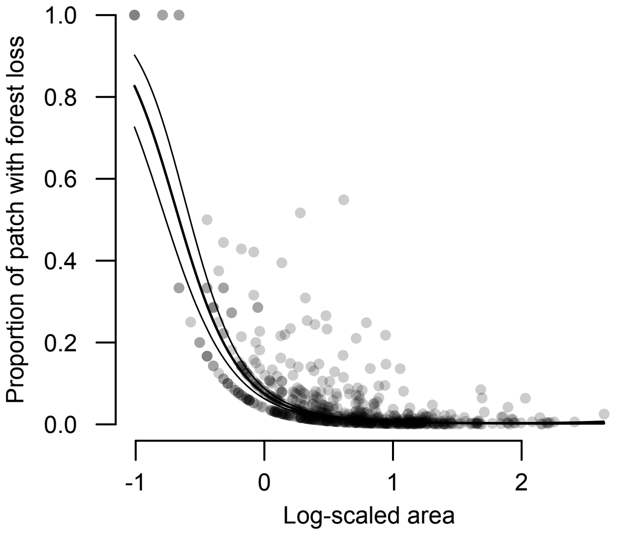


Figure S6. The effect of patch area on the proportion of forest loss in the marine transgression zone is a quadratic relationship. The black lines show the posterior mean and 95% credible intervals of the modeled relationship between area and forest loss when all other variables are held constant at their means. Each gray dot is a patch that experienced some loss.

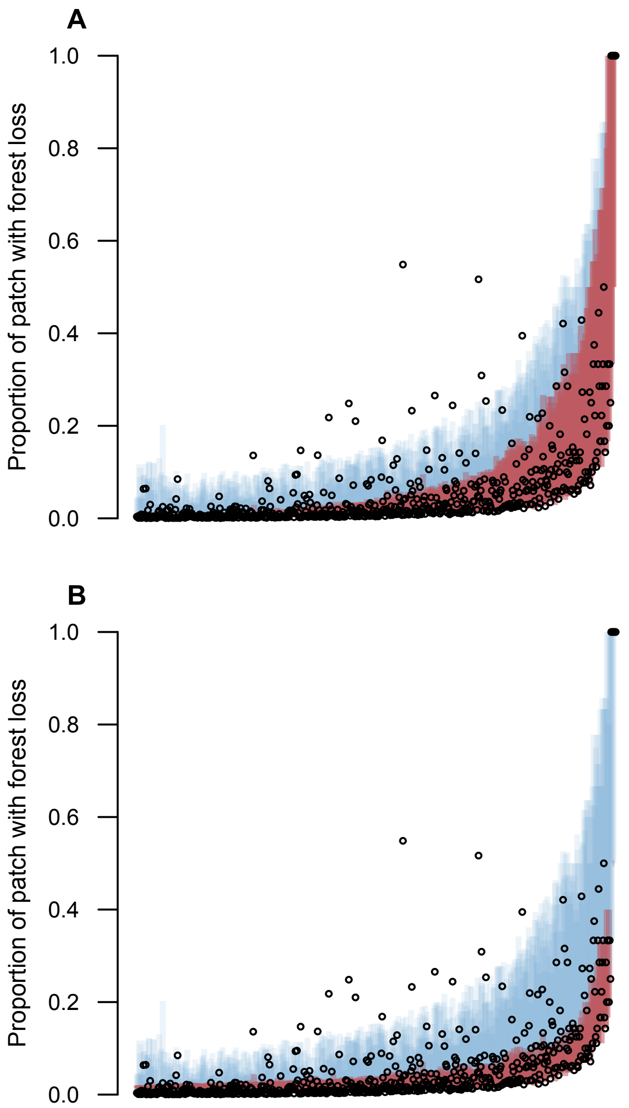


Figure S7. Posterior predictions compared to observed values for the model of the proportion of forest loss in the marine transgression zone. For both plots, posterior predictions are shown as vertical bars, ordered from low to high along the x-axis. Red bars show the 95% bounds for each prediction, not including sampling variation, and blue bars show the 95% bounds of the predictions when the sampling variation is included. Observed values for patches with at least some loss are shown as open circles. Plot **A** shows the posterior predictions for the full model of loss, which included variables for potential drivers, as described above; plot **B** shows the predictions for the same model without potential drivers. Both models contained ~ 95% of the values within the total predictive bounds (red plus blue), as expected, but the posterior predictions from the model that included drivers had greater predictive power: posterior predictions from the predictive component (red bars) of the model with drivers (**A**) contained the observed values more often predictions from the model without drivers (79% vs. 60%), especially for patches with more loss (further along the right side of the x-axis).

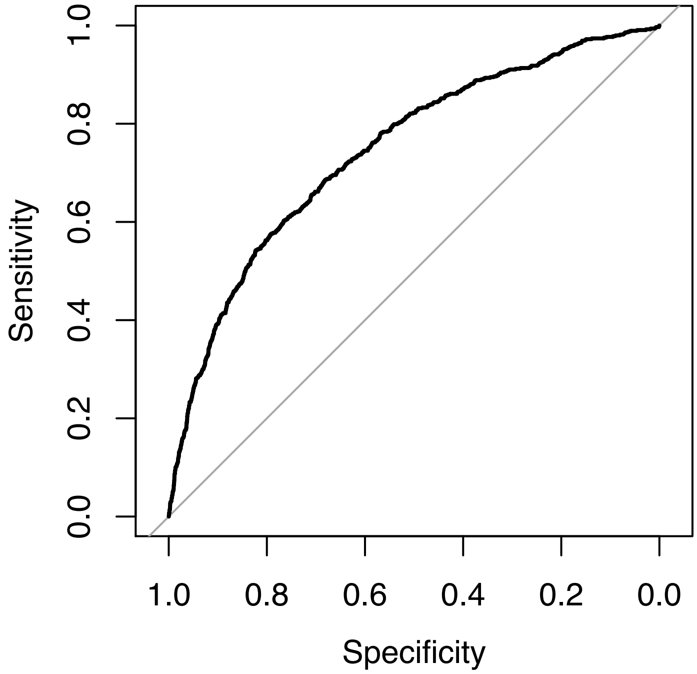


Figure S8. Receiver Operating Characteristic (ROC) curve for the model of the probability of at least some loss, showing the relationship between the true positive rate (sensitivity) and the true negative rate (specificity). The gray line shows the curve for a model with no predictive ability. The model's improvement over no predictive ability was consistent over the range of sensitivity and specificity values.

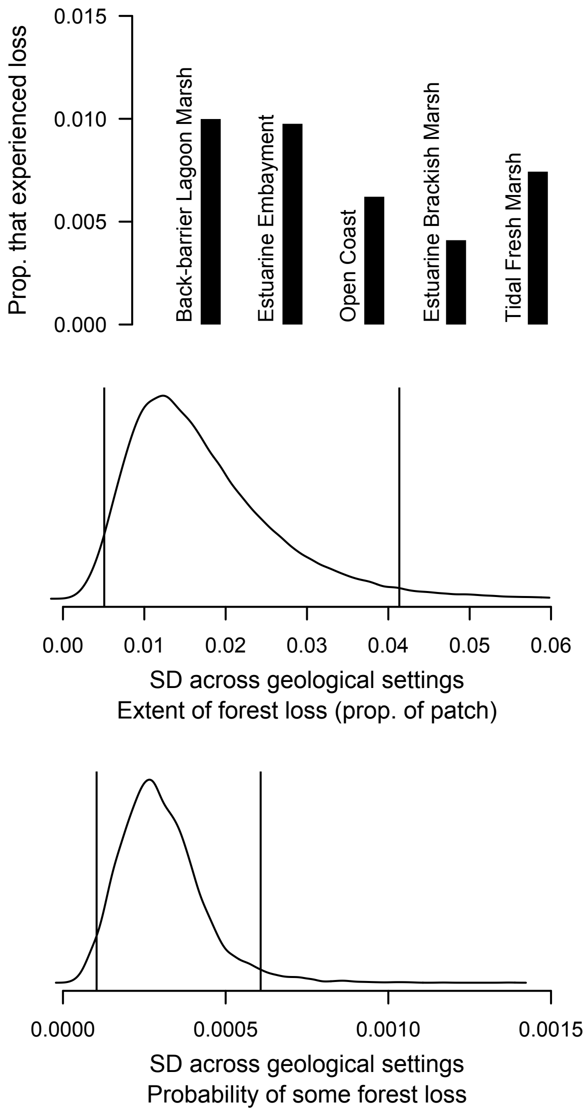


Figure S9. The influence of geomorphic setting on coastal forest loss. Top panel shows the proportion of coastal forest within 30 m of tidal marsh that experienced loss between 2000-2018 for five categories of geomorphic setting. The bottom two panels show the posterior estimate of the standard deviation of the categorical effects for geomorphic setting in the models of the extent of forest loss and probability of at least some loss. For both models, the bulk of the density is well above zero, suggesting that loss varies by geomorphic setting after accounting for the other biophysical variables in the model.

Table S1. Pearson's correlation coefficients of variables that potentially predict coastal forest loss.

| | **Latitude** | **Slope** | **Area** | **Prop. evergreen** | **SLR rate** | **Sandy storm surge** |
|---|---|---|---|---|---|---|
| **Latitude** | . | 0.50 | 0.09 | 0.29 | -0.86 | 0.44 |
| **Slope** | 0.50 | . | -0.08 | 0.25 | -0.42 | 0.13 |
| **Area** | 0.09 | -0.08 | . | 0.19 | -0.05 | 0.16 |
| **Prop. conifer** | 0.29 | 0.25 | 0.19 | . | -0.26 | -0.04 |
| **SLR rate** | -0.86 | -0.42 | -0.05 | -0.26 | . | -0.38 |
| **Sandy storm surge** | 0.44 | 0.13 | 0.16 | -0.04 | -0.38 | . |

**Supplementary code**

*Code for loading and manipulating Google Earth Engine data*

JavaScript code for loading, manipulating, and exporting the forest loss dataset and potential explanatory variables in Google Earth Engine (https://code.earthengine.google.com). Local datasets (peak storm surge during Hurricane Sandy) and datasets from the Earth Engine Data Catalog (Hansen et al. 2018, the National Land Cover Dataset, and a Digital Elevation Model) are loaded and masked by the extent of tree cover at the start of the time series (2000). A summary statistic from each dataset is calculated and exported for each multi-part polygon of the layer of coastal forest patches.

```
// load datasets
var patches = ee.FeatureCollection('');
var sandyVector = ee.FeatureCollection('');
var gfc2018 = ee.Image('UMD/hansen/global_forest_change_2018_v1_6');
var NLCD = ee.ImageCollection('USGS/NLCD');
var DEM = ee.Image('USGS/NED');

// select the band for landcover
var nlcd2011 = NLCD.filter(ee.Filter.eq('system:index', 'NLCD2011'));
var imgNLCD = ee.Image(nlcd2011.select(0).first());
var landcover = imgNLCD.select(['landcover']);
// get layer for natural cover (everything but bands 23 and 24; medium and hi
gh intensity development)
var natural = landcover.neq(23).and(landcover.neq(24));
// get layer for evergreen (42) or mixed cover (43)
var evergreen = landcover.eq(42).or(landcover.eq(43));
// get layer for any kind of forest cover (41 deciduous; 42 evergreen; 43 mix
ed)
var NLCDforest = landcover.eq(41).or(landcover.eq(42).or(landcover.eq(43)));

// get just the band for forest loss, turning the Hansen et al. collection in
to a single image
var lossImage = gfc2018.select(['loss']);
// get just the band for forest gain, turning the collection into a single im
age
var gainImage = gfc2018.select(['gain']);
// get just the band for tree cover in 2000, which gives total forest extent
at start of time series
var cover2000 = gfc2018.select(['treecover2000']);
// get just the band for the year of loss
```

```
var lossyear = gfc2018.select(['lossyear']);
// convert treecover 2000 into a binary image; only use pixels greater than 2
0% cover;
// same threshold as forest on NLCD
var cover2000Binary = cover2000.gt(20);
// mask loss and year layers by pixels that were forest in 2000 using 20% thr
eshold
var lossImageMask = lossImage.mask(cover2000Binary);
var gainImageMask = gainImage.mask(cover2000Binary);
var lossYearMask = lossyear.mask(cover2000Binary);
var canopyMask = canopy.mask(cover2000Binary);
// mask year of loss by pixels that experienced loss to get rid of zeros
var lossYearMaskNonzero = lossYearMask.mask(lossImageMask);
var lossSandy = lossYearMaskNonzero.gt(12)

// select the elevation band from DEM
var elevation = DEM.select('elevation');
// calculate slope
var slope = ee.Terrain.slope(elevation);
// mask slope by forest cover in 2000 using 20% threshold
var slopeMask = slope.mask(cover2000Binary);

// create an image for peak storm surge during Hurricane Sandy from the vecto
r file
var sandy = sandyVector.reduceToImage({properties: ['peak_storm'], reducer: e
e.Reducer.first()});

//reduce image by features for multiple polygons
var lossStats = lossImageMask.reduceRegions({
  reducer: ee.Reducer.sum().unweighted(),
  collection: patches,
  scale: 30
});

Export.table.toDrive({
  collection: lossStats,
  description: 'lossStats',
  fileFormat: 'CSV'
});
```

*Model code*

Code in base R and using JAGS pseudo-code to specify a simulation that generates known values and estimates them using a statistical model of the same structure as the primary model used to estimate the factors influencing forest loss. Two correlated independent variables are drawn from a multivariate normal distribution and then used in a regression equation to generate forest loss events. These variables are then centered and used in a statistical model of forest loss to demonstrate 1) that the effect of area on probability of loss can be distinguished from the fact that larger patches have more opportunities to experience loss, and 2) that a variable that is correlated with area can be estimated without bias.

```
# load required packages
library(R2jags)
library(R2WinBUGS)
# package MASS is used to generated correlated variables
library(MASS)

# generate correlated variables for area and latitude, with r = 0.10; for 500
0 data points
covs <- mvrnorm(5000, mu=c(30, 1000), Sigma=matrix(c(5*5, 0.1*5*1000, 0.1*5*1
000, 1000*1000), nrow=2), empirical=TRUE)
# if area is less than zero, replace with one
covs[covs[ ,2] < 1, 2] <- 1
# round area to simulate number of pixels to allow use in the binomial
covs[,2] <- round(covs[,2])
# check correlation to make sure it is ~0.10
cor.test(covs[,1], covs[,2])

# SD of overdispersion
sd <- 0.25
# overdispersion for 5000 patches
rand <- rnorm(5000, 0, sd)
# N is area (number of pixels)
N <- covs[,2]
lat <- covs[,1]
# probability of loss (Pr of zero loss is 1 - q)
q <- 0.00025
# standardize and center N and latitude
N_cov <- (log(N) - mean(log(N)))/(2*sd(log(N)))
lat_cov <- (log(lat) - mean(log(lat)))/(2*sd(log(lat)))
# generate realized values for whether a patch experienced loss
index <- rbinom(5000, N, q)
index[index>0] <- 1
# simulated logistic regression equation to generate known values
mu <- -4 + rand - 0.5*lat_cov - 1*N_cov
# back-transform logistic regression equation
p <- exp(mu)/(1+exp(mu))
# generate data using area (N) and regression equation
data_temp <- rbinom(5000, N, p)
# get data with zeros from inflation component
data <- data_temp*index
```

```
# create data vectors for JAGS for patches that have at least some loss
dataJAGS <- data[index>0]
Njags <- N[index>0]
N_covJAGS <- N_cov[index>0]
lat_covJAGS <- lat_cov[index>0]

JAGS <- function(){
  mu ~ dnorm(0, 0.04)
  B ~ dnorm(0, 0.04)
  C ~ dnorm(0, 0.04)
  sd ~ dunif(0, 10)
  tau <- 1/(sd*sd)
  for(i in 1:length(dataJAGS)){
    int[i] ~ dnorm(0, tau)
    logit(p[i]) <- mu + C*lat_covJAGS[i] + int[i] + B*N_covJAGS[i]
    dataJAGS[i] ~ dbin(p[i], Njags[i]); T(1,)
  }
}

if (is.R()){
  filename <- file.path(tempdir(), "JAGS.bug")}
write.model(JAGS, filename)
inits <- list(list(B=1, C=0))
data <- list("dataJAGS", "Njags", "N_covJAGS", "lat_covJAGS")
parameters <- c("p", "mu", "B", "int", "sd", "C")
JAGS <- jags(data=data, inits=inits, parameters.to.save=parameters, filename,
              n.chains=1, n.burnin=100000, n.iter=100000, n.thin=1, DIC=TRUE)
```

Just Another Gibbs Sampler (JAGS) pseudo-code for estimating the probability that a patch experiences at least some forest loss. Forest loss is specified as the inverse (patch experiences loss is 0; no loss is 1) because the binomial probability, q, is specified as the probability of no pixels in the patch experiencing loss. The probability of loss, p, from the regression equation is transformed to q before being fed to the binomial sampling distribution. Priors for two alternative specifications, more and less prior information, are shown.

```
# prior for parameters that describes the SD and precision of regression coef
ficients
sdC ~ dunif(0, 100)
# alternative prior that specifies more prior information
#sdC ~ dunif(0, 10)
tauC <- 1/(sdC*sdC)
# prior for intercept of regression equation
mu ~ dnorm(0, 0.04)
```

```
# alternative prior that specifies more prior information
#mu ~ dnorm(0, 0.25)
# regression coefficients are drawn from a t distribution with 4 degrees of f
reedom
for(e in 1:8){
  C[e] ~ dt(0, tauC, 4)
}
# regression equation that includes variables that potentially explain the pr
obability of at least some forest loss
for(i in 1:length(loss_bin_inv)){
  logit(pi[i]) <- mu + C[1]*slope[i] + C[2]*area[i] + C[3]*area[i]*area[i] +
C[4]*evergreenProp[i] + C[5]*SLR[i]
  + C[6]*sandy[i] + C[7]*slope[i]*SLR[i] + C[8]*evergreenProp[i]*lat[i]
  # p is the probability of a pixel in a given patch experiencing loss; speci
fy, q, the probability of no pixels in a patch experiencing loss
  q[i] <- (1-pi[i])^cover[i]
  # whether a patch experienced loss (0) vs. no loss (1) is a binomial variab
le with probability q
  loss_bin_inv[i] ~ dbern(q[i])
  }
```

Just Another Gibbs Sampler (JAGS) pseudo-code for estimating the probability of a pixel in a patch experiencing loss, give at least some loss at the patch-scale, which is equivalent to the proportion of the patch that experienced loss. Only data from patches that experienced loss in at least one pixel are used. Priors for two alternative specifications, more and less prior information, are shown.

```
sd_int ~ dnorm(0, 0.04)
#sd_int ~ dnorm(0, 0.25)
# prior for parameters that describes the SD and precision of regression coef
ficients
sdB ~ dunif(0, 100)
# alternative prior that specifies more prior information
#sdB ~ dunif(0, 10)
tauB <- 1/(sdB*sdB)
# prior for intercept of regression equation
mu ~ dnorm(0, 0.04)
# alternative prior that specifies more prior information
#mu ~ dnorm(0, 0.25)
# regression coefficients are drawn from a t distribution with 4 degrees of f
reedom
for(e in 1:10){
  B[e] ~ dt(0, tauB, 4)
  }
# regression equation that includes variables that potentially explain the pr
```

```
obability of at least some forest loss
for(i in 1:length(loss_cond)){
  # variation in the SD of the random effect is explained by coefficients for
slope and area
  log(sd[i]) <- sd_int + B[9]*slope_cond[i] + B[10]*area_cond[i]
  tau[i] <- 1/(sd[i]*sd[i])
  # overdispersion relative to the binomial distribution is explained by a no
rmal distribution
  overdisp[i] ~ dnorm(0, tau[i])
  # regression equation that includes variables that potentially explain the
probability of forest loss, given that there was at least some
  logit(p[i]) <- mu + overdisp[i] + B[1]*slope_cond[i] + B[2]*area_cond[i] +
B[3]*area_cond[i]*area_cond[i]
  + B[4]*evergreenProp_cond[i] + B[5]*SLR_cond[i] + B[6]*sandy_cond[i] + B[7]
*slope_cond[i]*SLR_cond[i]
  + B[8]*evergreenProp_cond[i]*lat_cond[i]
  # the number of pixels that experienced forest loss is a binomial variable
with probability p and size equal to the number of pixels that are forest
  loss_total[i] ~ dbin(p[i], cover_cond[i]); T(1,) # the binomial distributio
n is truncated to exclude the possibility of zeros
  }
```

Just Another Gibbs Sampler (JAGS) pseudo-code for estimating trends in the rate of forest loss by latitudinal bands (each band is 0.5 decimal degrees). Baseline rates of loss and trends for each band are both drawn from a distribution that has a global mean, which is the trend in the rate of forest loss for the entire study area. Each band is specified to have an independent parameter for sampling variation. The same model structure is used to estimate for each year the number of patches in each band experienced forest loss (data specified as “events_binned” in the code below).

```
# prior for the mean trend in forest loss over time
Bmu ~ dnorm(0, 0.0001)
# prior for the mean baseline rate of forest loss
int_mu ~ dnorm(0, 0.0001)
# prior for the SD and precision of the band-level trends in forest loss over
time
Bsd ~ dunif(0, 1000)
Btau <- 1/(Bsd*Bsd)
# prior for the SD and precision of the band-level baseline rates of forest l
oss
int_sd ~ dunif(0, 1000)
int_tau <- 1/(int_sd*int_sd)
```

```
# for each band of latitude
for(z in 1:length(loss_binned[,1])){
  # baseline rate of forest loss for each band, int, is drawn from a global m
ean, int_mu
  int[z] ~ dnorm(int_mu, int_tau)
  # trend in forest loss over time for each band, B, is drawn from a global m
ean, B_mu
  B[z] ~ dnorm(Bmu, Btau)
  # each band is allowed to have a separate parameter for sampling variation
  sd[z] ~ dunif(0, 1000)
  tau[z] <- 1/(sd[z]*sd[z])
  # for each year, i
  for(i in 1:length(loss_binned[1,])){
    # regression equation that includes the potential for trends in the likel
ihood of forest loss over time
    mu[z, i] <- int[z] + B[z]*i
    # the number of pixels with by latitude band and year is normally distrib
uted
    loss_binned[z, i] ~ dnorm(mu[z, i], tau[z])
    # use the model for the number of patches with new loss in each year
    #events_binned[z, i] ~ dnorm(mu[z, i], tau[z])
    }
  }
```
